# A dermal-epidermal junction-inclusive skin model enabled by controllable hydrogel swelling

**DOI:** 10.64898/2026.06.29.735406

**Authors:** Tobias Hammer, Tobias Spirig, Markus Rottmar, Katharina Maniura-Weber, Kongchang Wei, René M. Rossi

## Abstract

Tissue engineered skin models are important tools for the *in vitro* study of physiological and pathophysiological processes as well as the evaluation of therapeutic strategies and the efficacy of pharmaceutical and cosmetic compounds. Replicating the functional anatomy of cutaneous tissue is a crucial aspect in ensuring that observations made using these models are translatable to the actual situation in native skin. However, most contemporary full-thickness skin models neglect the reconstruction of the undulated microtopography of the dermal-epidermal junction (DEJ), which not only contributes to the biological functionality of the skin (e.g. stem cell niches), but also affects tissue mechanics and drug diffusion. Herein, we fabricated bilayer skin models with DEJ-like microtopographies introduced by interfacial wrinkling between a hydrogel and a nanofibrous membrane through a controllable swelling-deswelling approach. The interfacial wrinkles contributed to the structural integrity of the bilayer models. Their formation could be induced in the presence of living cells through mechanical stress-driven buckling instabilities, thus differentiating the process from commonly used pre-patterning techniques. Bilayer models supported the co-culture of human dermal fibroblasts and human epidermal keratinocytes, and the formation of stratified epithelia. Our findings provide a potential alternative method to introduce DEJ-like anatomical features into full-thickness skin tissue models.

## Introduction

As one of the largest and most dynamic organs of the human body that serves as both a semi-permeable barrier against as well as a major interface for engaging with the exterior environment. The skin performs several different functions pertaining to physiological homeostasis and pathological defense. While the inherent regenerative properties of skin allow it to easily repair small superficial damages, larger and deeper wounds usually require medical intervention to ensure appropriate healing [1, 2]. Since the advent of tissue engineering, and especially over the past years, there was great interest in fabricating artificial skin constructs for the purpose of medical grafting [3], evaluation of wound healing therapies [4, 5], modeling of biological processes [6, 7], or screening of pharmaceutical and cosmetic compounds [8]. However, these constructs are often limited in their biological and anatomical complexity. While most tissue engineered skin constructs focus on reconstructing the outermost epidermis and the underlying dermis, the undulated intersection between these layers, called the dermal-epidermal junction (DEJ), is largely neglected [9].

The DEJ exhibits a distinctive topography that is characterized by the presence of epidermal rete ridges and dermal papillae, which form fingerlike interdigitations with dimensions ranging from 50-400 μm in width and 50-200 μm in depth [10]. These features play a crucial role in the overall structural integrity of the tissue and its mechanical properties by increasing the effective surface area between the two layers, thus allowing for a greater number of hemidesmosomes and enhanced cellular adhesion [11]. Furthermore, rete ridges act as niches for epidermal stem cells, affecting both cellular proliferation and differentiation as well as wound healing and tissue homeostasis [12]. Thus, it becomes apparent that the DEJ constitutes a crucial element of the functional anatomy of skin tissue, the significance of which is further supported by the dynamic changes to its morphology due to ageing, scarring, or pathological progression (e.g. epidermolysis bullosa, psoriasis, cancer, etc.) and the associated increased susceptibility to shear stress, blister formation and loss of structural integrity [13].

Despite its importance in terms of tissue functionality and interaction with external stimuli (e.g. drug absorption, paracrine signaling, mechanical stress, etc.), most contemporary tissue engineered constructs neglect to recreate the DEJ in full-thickness skin models. Instead, models opt for a plane interface between the epidermal and dermal compartments, which results in a homogeneous distribution of cells and biomolecules that is significantly different from the heterogeneous arrangement of natural skin [14–16]. As the influence of mechanical and structural stimuli in directing biological events and tissue function has garnered increased recognition in recent years, more focus has been directed at mimicking the biophysical and anatomical features of target tissues [17, 18]. Consequently, studies that aim at implementing the DEJ into tissue engineered skin constructs have become more common. The reported approaches generally rely on imparting a DEJ-like topography onto a static surface through bioprinting [19–21], photolithography [22–25], laser ablation [26, 27], or electrospinning [19, 21, 23]. Insights gained from observations of the fetal development of the DEJ as well as computational simulations of the biomechanical forces acting on epidermal and dermal layers, however, point toward the involvement of mechanical stresses and subsequent buckling instabilities in the formation of papillary networks and rete ridges [28–30]. Differential volumetric growth and geometrical constraints between the soft dermal layer and the stiffer epidermal layer are suggested to be the main driving force behind this phenomenon [29, 30]. Furthermore, it has been shown that the undulated morphology of the DEJ causes persistent heterogeneities in local stresses along its interface, which could affect cellular behavior and niche formation through mechanical stimulation [29].

Exerting precise control over the volumetric changes in living matter represents a challenging endeavor, as they are often the result of complex cellular and metabolic pathways leading to the synthesis or degradation of tissue. Tissue engineered scaffolds fabricated from functionalized synthetic or biological polymers on the other hand are more amendable to precise manipulation due to the inclusion of chemical modifications that endow these materials with specific properties. Biopolymers like gelatin or collagen are particularly promising components for these scaffolds, since their inherent bioactive properties and biodegradability allow them to directly interact with and be modulated by incorporated cells [31]. Recently, we developed a hydrogel formulation based on the chemical modification of cold-water fish gelatin with thiol and norbornene groups that allows for the fabrication of hydrogels with controllable swelling behavior [32]. As both the swelling and deswelling process can be conducted under cytocompatible conditions, we are able to perform dynamic volumetric alterations that enable the formation of stress-driven buckling instabilities and mechanical stimulation in the presence of cells.

Here we describe the formation of a full-thickness skin tissue construct exhibiting a wrinkled interface between the dermal and epidermal layers facilitated by swelling-deswelling-mediated volumetric changes of a cold-water fish gelatin-based hydrogel (cfGel-Hydrogel). Wrinkle formation is driven by interfacial stress generated between a soft swollen hydrogel and a stiffer electrospun membrane by triggering volumetric shrinkage of the former, resulting in buckling instabilities due to geometrical constraints caused by the adhered membrane. Co-culturing of human dermal fibroblast (HDFs) and human epidermal keratinocytes (HEKs) in this skin-mimicking scaffold lead to the formation of a DEJ-inclusive tissue engineered skin model with architectural features that represent a closer approximation of the native anatomy of skin, thus making the model a potential future candidate to be used for the study of skin-related physiological and pathophysiological conditions or pharmacochemical testing.

## Results and discussion

### Design of skin-mimicking bilayer scaffolds

Scaffolds were comprised of two distinct elements, namely a soft hydrogel base and an electrospun nanofibrous membrane. Hydrogels, which were based on our previously developed cfGel-Hydrogel formulation [32], acted as a dermal mimic owing to their structural resemblance to cutaneous ECM and amenability for the direct encapsulation of HDFs during photopolymerization. Hydrogels were fabricated using custom-made PDMS/glass molds with variable dimensions. Nanofibrous membranes were electrospun from a solution of unmodified cold-water fish gelatin and D-fructose in a controlled environment (25 °C, 46% relative humidity) according to previously published parameters found in literature with slight variations [33, 34]. Membranes were subsequently crosslinked at 140 °C for 8 h. Herein, we henceforth refer to the crosslinked membranes as “cfGel-Membranes”. The nanofibrous topography of cfGel-Membranes mimics the fibrous structure of the natural basement membrane that separates the epidermis from the underlying dermis, thus acting as a growth substrate for primary HEKs. The small pore size of nanofibrous cfGel-Membranes also prevents the infiltration of both fibroblasts and keratinocytes into the epidermal and dermal layers respectively, keeping the two cell populations separated. Nanofibers exhibited a straight morphology and smooth surface structure as well as a random orientation (**Fig. 1A**), with an average fiber diameter of 215.42 ± 32.07 nm (**Fig. 1B**).

**Figure 1.**
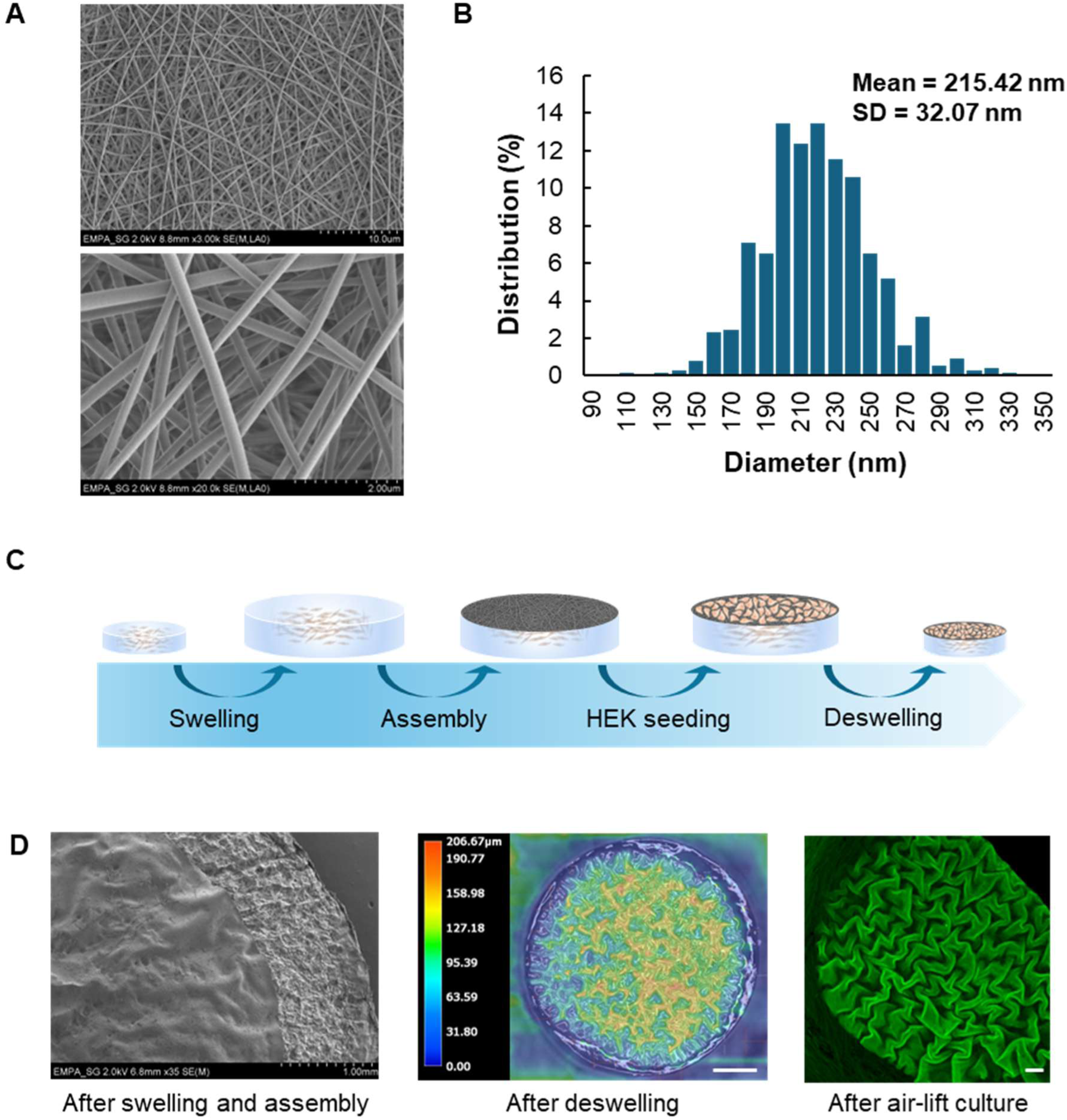
Bilayer scaffold components and assembly. (**A**) SEM images of electrospun cold-water fish gelatin membranes. (**B**) Average fiber diameter of nanofibrous membranes. (**C**) Schematic of multistep assembly process used for bilayer scaffolds. (**D**) SEM, color-coded optical microscope profilometry, and confocal microscopy images depicting the surface topography of bilayer scaffolds at various timepoints. SEM image depicts a section of the freeze-dried bilayer with the electrospun membrane on top of the swollen cfGel-Hydrogel. Green fluorescence of nanofibrous membranes in confocal images is facilitated by the inclusion of fluorescently labeled dextran (FITC-dextran). Left, middle: Scale bar = 1 mm, right: Scale bar = 200 μm.

Assembly of bilayer scaffolds followed a multi-step process (**Fig. 1C**). Initially, HDF-inclusive cfGel-Hydrogels were fabricated as described in our previous study [32] and subjected to controlled swelling for 1 h at 37 °C in an isotonic sucrose solution (0.25 M). Afterwards, dry cfGel-Membranes were placed on top of the swollen hydrogels, followed by seeding of HEKs and subsequent deswelling of the constructs in cell culture media. Mounting interfacial stresses between the shrinking cfGel-Hydrogels and the cfGel-Membranes induced by the deswelling process resulted in buckling instabilities that facilitated the formation of wrinkles (**Fig. 1D**).

### Wrinkle formation mediated by swelling-deswelling-induced volumetric changes

Given that volumetric changes in cfGel-Hydrogels are the driving force of wrinkle formation in our system, we first assessed how the degree of swelling and deswelling time affects wrinkle dimensions. For this, we subjected cfGel-Hydrogels to varying pre-swelling times in an isotonic solution of sucrose (0.25 M) and recorded the changes in both size and weight before subsequent deswelling in PBS (**Fig. 2A**) [35]. We chose isotonic sucrose as the swelling medium since it can be used to swell cfGel-Hydrogels without causing cell lysis due to hypotonic conditions, thus allowing for direct encapsulation of cells in our dermis-mimicking layer and the initiation of volumetric changes at a later stage, which is crucial for our intended co-culture application. In an earlier study, we verified that the swelling behavior of cfGel-Hydrogels in isotonic sucrose solutions is identical to ultrapure water (UPW), which allows translation of results from swelling-deswelling-related measurements between the two media systems [32]. According to our results, cfGel-Hydrogels experience the most pronounced increase in diameter within the first 30 min of incubation. After 15 min, diameters increased in size from 6 mm to 8.15 ± 0.12 mm, reaching 9.96 ± 0.22 mm after a total swelling time of 30 min. Subsequent increases were less prominent with 10.40 ± 0.26 mm after 45 min and 10.73 ± 0.25 mm after 60 min of swelling. Interestingly, following exposure to PBS for 30 min, all samples reverted close to their original dimensions, regardless of prior swelling time, with average diameters of 6.33 ± 0.09 mm. Prolonged exposure to PBS for up to 24 h resulted in further deswelling of samples, with diameters decreasing to an average of 5.79 ± 0.03 mm (**Fig. 2B**).

**Figure 2.**
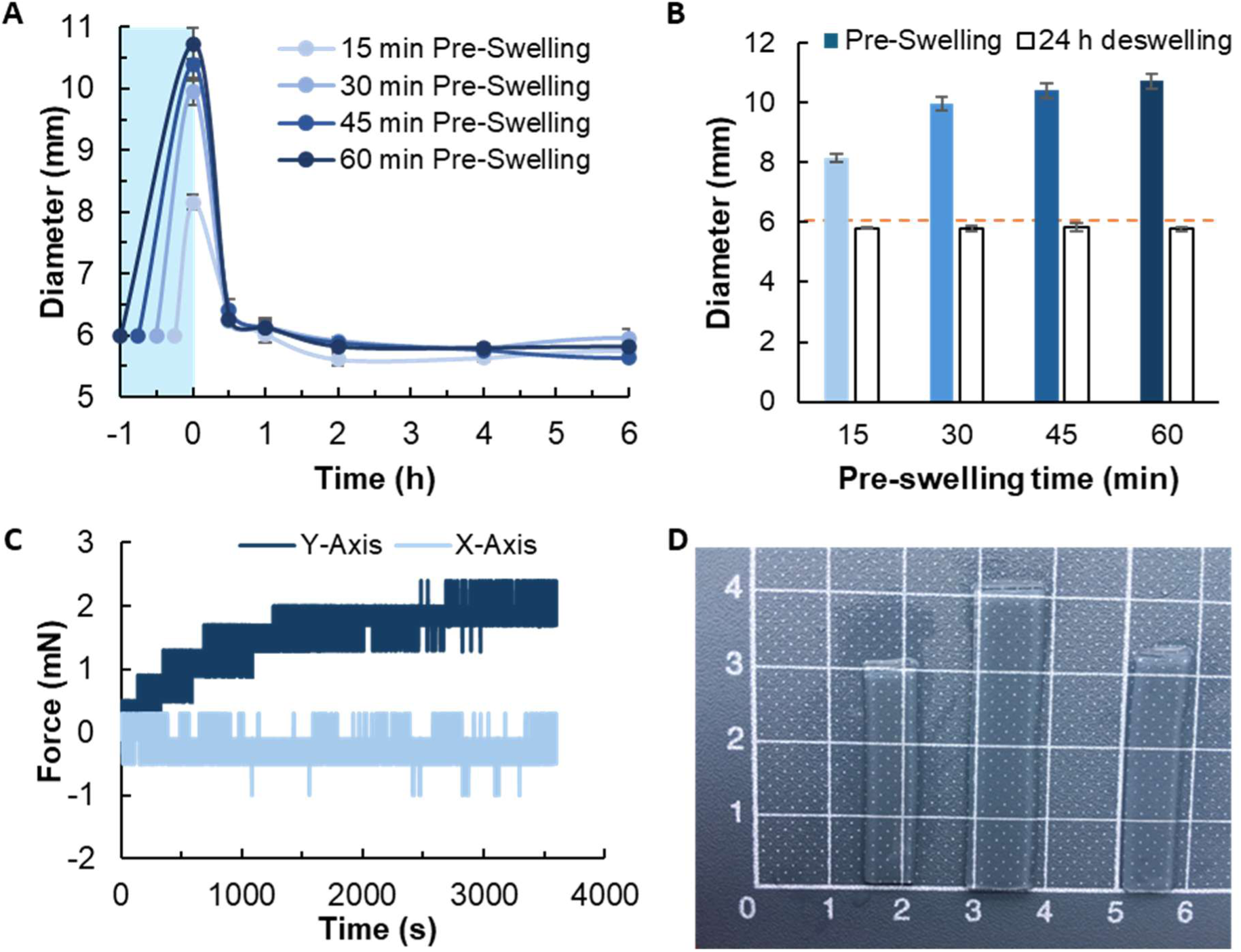
Swelling-deswelling-induced volumetric changes and force generation. (**A**) Evaluation of volumetric expansion of cfGel-Hydrogels as a function of pre-swelling time. Blue area indicates pre-swelling in isotonic sucrose solution prior to deswelling in PBS at t = 0 h. *n = 3* (**B**) Diameters of cfGel-Hydrogels after swelling for 15, 30, 45, and 60 min. After deswelling for 24 h, all samples displayed similar diameters close to their original dimensions (indicated by dashed line) irrespective of prior swelling. Error bars indicate SD. *n = 3* (**C**) Representative force curve generated by cfGel-Hydrogels during deswelling in PBS over the course of 1h. Y-Axis displays force generated by shrinking samples that were pre-swelled for 1h, while X-Axis displays force measured by an empty load cell as internal reference. (**D**) Morphology of samples used for force-detection measurement. From left to right: before swelling; after swelling; after deswelling. The marked grid distances correspond to 1 cm.

Employing a swelling-deswelling-based approach to facilitate volumetric changes in cfGel-Hydrogels has several advantages over mechanical stretch-release-reliant methods for interfacial wrinkle formation [36–38]. Firstly, swelling enables a isometric expansion of samples, resulting in a homogeneous distribution of force, which can be challenging to achieve through alternative methods, especially in the presence of living cells. Swelling also results in expansion along the z-direction, whereas mechanical stretch typically results in a concomitant thinning of hydrogels, which might affect the morphology of interfacial wrinkles in our bilayer system. Mechanical stretching of hydrogels generally requires their prior fixation with clamps, hooks, or other devices at multiple points, depending on the direction of deformation, resulting in localized differences to both volumetric changes and experienced forces between the fixed and non-fixed parts. Applying uniform mechanical stretch along multiple directions can therefore be a challenging endeavor that requires complex setups and devices with high precision over exerted forces. For hydrogels with intricate morphologies, small sizes, or restrictive environments (e.g. cell culture conditions) such an approach might not be feasible in the first place. Compared to the instantaneous reversion enabled by stretch-release, the speed of swelling-deswelling-mediated volumetric changes is much slower. In our case, cfGel-Hydrogels exhibit a significant increase in size after as little as 15 min of swelling while also being able to revert close to their original dimensions following exposure to PBS for approximately 30 min. A more gentle and slower volumetric alteration can be beneficial for applications that involve cell-laden hydrogels, as it gives cells more time to respond to the changing environment.

The compressive forces exerted on cfGel-Membranes by shrinking cfGel-Hydrogels ultimately leads to the former’s deformation and the emergence of surface wrinkles. As such, we attempted to evaluate the amount of force generated during the deswelling process of cfGel-Hydrogels. To this end, swollen hydrogels of rectangular shape were fixed onto a biaxial test machine and subsequently deswelled in PBS (Supporting Information, **Fig. S1**). Hydrogels were fixed along the Y-axis while the load cells along the X-axis were left empty to detect and account for any passive drift in force during the measurement (**Fig. 2C**). The results indicate a steep increase in force within the first ∼1000 s of the measurement, with curves eventually plateauing around 2.12 ± 0.61 mN. Interestingly, force values remained persistent over the entire duration of the measurement without any sign of decrease, implying that no dissipation of residual stress through viscoelastic stress-relaxation occurred after 1 h. Releasing the fixed samples from the load cells resulted in a complete reversion to their original dimensions prior to swelling, further suggesting the absence of notable permanent viscoelastic deformations (**Fig. 2D**).

Numerical simulation experiments using 2D multilayered skin models have placed emphasis on the influence of residual mechanical stresses in the development of the wrinkled interface during rete ridge formation in native skin tissue [30]. Deformation of the epidermal layer results in the formation of undulations that exhibit localized variations in perceived levels of compression. For example, valleys on the dermal side experience only mild compressive forces, while epidermally located valleys and peaks are under higher compression that limits their expansion [30]. This heterogeneous distribution of compressive forces along the DEJ-interface results in cells positioned at different locations to experience variable levels of mechanical stimulation, thus altering their behavior in accordance [39]. Considering the importance of mechanical signaling in the dynamic modulation of cellular behavior and proper tissue growth, it is likely that simply mimicking the topographical structure of the DEJ through contemporary patterning techniques is insufficient in inducing native cell responses [9]. Instead, hydrogel deswelling-induced mechanical stress can both lead to the formation of DEJ-like topographical features and expose the seeded keratinocytes to localized variations in compressive force during wrinkle formation. The gradual deformation of the DEJ-mimicking cfGel-Membranes at a slower rate compared to mechanical stretch-release allows the seeded keratinocytes to experience the spatiotemporal shifts in their growth environment, as opposed to being directly seeded onto a pre-patterned surface. During the development of the epidermis, exposure of keratinocytes to variable mechanical cues according to the surrounding DEJ-like micro-topographies could potentially regulate their proliferation and differentiation in a similar manner to native rete ridges and dermal papillae. This feature makes swelling-deswelling a promising approach in developing scaffolds with included architectural and mechanical stimuli with potential similarity to native skin. However, further studies relating to the influence of mechanical and structural stimuli on epidermal development will have to be conducted in the future in order to draw definitive conclusions regarding their importance.

### Controlled formation of surface wrinkles through adjusted pre-swelling

In the next step, we evaluated how swelling time correlates with the formation of surface wrinkles. To this end, cfGel-Hydrogels pre-swelled for 0 min, 15 min, 30 min, 45 min and 60 min were fitted with a dry piece of cfGel-Membrane before deswelling overnight. The surface wrinkles were analyzed in order to obtain both surface profiles as well as cross-section images (**Fig. 3**). The surface height profile was reconstructed by optical focus-variation scanning and presented as a color-coded map (**Fig. 3A**). In the generated height map, red and yellow areas correspond to elevated surface regions, whereas blue and green areas indicate depressions. The color-coded profile reveals the formation of randomly oriented surface wrinkles with no preferred direction, indicating globally homogeneous deswelling of the cfGel-Hydrogel base. To quantify the surface height variation observed in the color-coded map, a line profile was extracted across the scanned area (**Fig. 3B-C**). We analyzed the surface wrinkles by measuring the depth and number of the formed valleys, as these would correspond to rete ridges in native skin as opposed to the peaks, which would be representative of dermal papillae. We found that valley depth did not significantly differ between different pre-swelling times (**Fig. 3D-E**). The numbers measured for samples with 15 min and 30 min of pre-swelling time were 27.9 ± 15.8 µm and 27.0 ± 13.8 µm respectively. Increasing pre-swelling time to 45 min and 60 min resulted in valleys with an average depth of 22.9 ± 10.8 µm and 23.4 ± 15.7 µm respectively. Unexpectedly, even samples with 0 min of pre-swelling time exhibited wrinkled surface profiles, albeit not as prominent, with valleys averaging depths of 19.5 ± 9.3 µm. A possible explanation for the presence of surface wrinkles even in the absence of pre-swelling could be a slight drying out and shrinkage of the hydrogel surface following attachment of the dry cfGel-Membrane due to the latter absorbing a small amount of hydrogel-bound liquid. This transfer of fluid from a wet substrate surface to a dry membrane has been reported to effectively promote interfacial interactions [40]. Previous studies have also demonstrated the impact of other factors such as membrane strain state, interfacial shear stress, isotropic/anisotropic loading, and the thickness of both substrate and film on the formation and evolution of surface wrinkles on bilayer soft materials [41, 42]. Therefore, the similarities in wrinkle profiles obtained with different pre-swelling times might be partially attributable to the combined effect of deswelling, liquid transfer, and constant thickness of the cfGel-Membranes [43]. Arithmetic mean height (Sa) of surface profiles obtained at different Gaussian filter cut-off frequencies (*λ*_c_) was also used as a statistical measure of average height deviation (**Fig. 3F** and Supporting Information **Fig. S2**). Varying the Gaussian filter cut-off allowed separation of large-scale and fine-scale surface features, thereby enabling a multi-scale characterization of the surface morphology. The results show no notable influence of *λ*_c_ on the overall trend of the different surface roughness parameters based on pre-swelling time.

**Figure 3.**
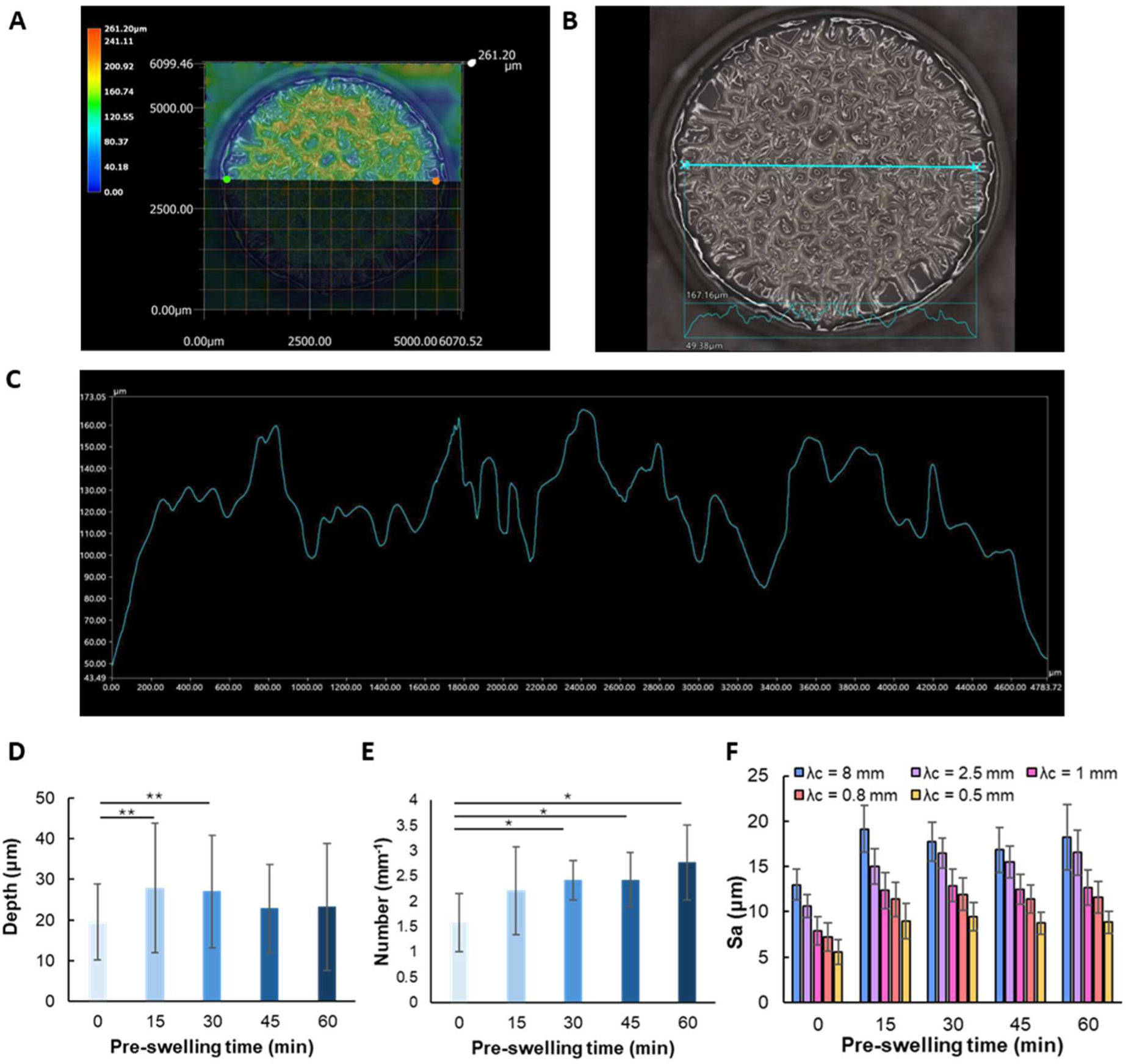
Effect of pre-swelling time on surface wrinkle formation. (**A**) Representative color-coded height map of surface profile. (**B**) Cross-sectioning of sample with 1 h of pre-swelling time. Line indicates the location and direction of the cross-section profile. (**C**) Cross-section height profile obtained from panel B. (**D**) Average depth of formed wrinkles for samples with different pre-swelling times. Error bars indicate SD, p-values were determined using a Kruskal-Wallis and Mann-Whitney U-test, p**≤ 0.01. *n = 6* (**E**) Average number of wrinkles for samples with different pre-swelling times. Error bars indicate SD, p-values were determined using a Student’s t-test, p**≤ 0.01, p*≤ 0.05. *n = 6* (**F**) Arithmetic mean height (Sa) of surface profiles obtained for samples with variable pre-swelling time at different Gaussian filter cut-off frequencies (λc).

According to literature, rete ridge dimensions range from 50-400 µm in width to 50-200 µm in depth, exhibiting differences across age, anatomical location and phototype, and being further influenced by pathological conditions [13, 27, 44]. Based on various studies that include data on the height of epidermal rete ridges across multiple factors (e.g. age, body location, pathophysiological conditions), our measured wrinkles exhibit most similarities to rete ridges situated in the facial and forearm regions [13]. However, the density of observed wrinkles in our samples with numbers averaging 1.58 ± 0.58 mm^−1^ for 0 min, 2.21 ± 0.87 mm^−1^ for 15 min, 2.42 ± 0.39 mm^−1^ for 30 min, 2.43 ± 0.53 mm^−1^ for 45 min, and 2.77 ± 0.74 mm^−1^ for 60 min (**Fig. 3E**), were significantly lower compared to what has been previously reported for epidermal rete ridges (e.g. young buttock/forearm: 10 ± 1 mm^−1^/6 ± 2 mm^−1^; aged buttock/forearm: 10 ± 3 mm^−1^/5 ± 3 mm^−1^) [45]. While most studies describe a flattening and concomitant reduction of dermal papillae, and consequently rete ridges, with age [46–48], some reports do not observe significant differences in rete ridge numbers in skin of young and old individuals [45]. Defining the exact number and dimensions of rete ridges and dermal papillae can be challenging, as their irregular shape can make them prone to variable interpretation. It should be noted that the absolute wrinkle dimensions obtained in this system are not intended to replicate the full geometric spectrum of native DEJ. Instead, the relevance of the model lies in reproducing key architectural features and mechanically generated interfacial undulations under cell-compatible conditions, rather than exact geometric equivalence. The wrinkle depths and densities observed here fall within the lower range reported for facial, forearm, and aged skin, which are characterized by shallower and less densely packed rete ridges. Accordingly, the present model captures a subset of physiologically relevant DEJ architectures rather than a single canonical geometry.

### Influence of surface wrinkles on bilayer adhesion

Apart from its topography-altering effects, the formation of surface wrinkles has also been shown to promote interfacial adhesion between various surfaces [49–53]. The ability to utilize dynamic alterations in surface patterns to establish adhesion is a trait often observed in natural systems and has been frequently exploited in the creation of surfaces with tunable physical and biological properties [49, 54, 55]. In this study, we evaluated whether the formation of interfacial wrinkles promotes the adhesion between cfGel-Hydrogels and cfGel-Membranes and to what extent. For this, we conducted shear and peel tests on wrinkled and non-wrinkled bilayer models to measure the required force to separate cfGel-Membranes from cfGel-Hydrogels both in air and in a submerged state (**Fig. 4A**). For shear tests, samples demonstrated similar curve shapes in force development irrespective of their state of submersion (**Fig. 4B-C**). In general, the detachment of wrinkled membranes required a lower maximum force compared to non-wrinkled membranes, but the force needed to be applied over a larger displacement range to obtain complete removal of membranes from hydrogels. Conversely, non-wrinkled membranes already exhibited complete detachment after approximately half the amount of displacement needed to completely detach wrinkled membranes, albeit with measured maximum forces typically exceeding the ones detected for the latter by more than double. A possible explanation for the observed behavior could be the formation of multiple smaller points of attachment across the entire bilayer interface due to wrinkle formation. While this results in relatively weak forces being required to detach individual contact points, their large number and distribution across the entire interface provides stability to the bilayer. Consequently, the range of displacement needs to cover the entire length of the membrane-hydrogel interface. This explanation is further supported by force curves for wrinkled samples frequently displaying a jagged morphology, indicating several smaller spikes in force during displacement. In the case of non-wrinkled interfaces, the contact area between membrane and hydrogel is larger, thus requiring greater force to facilitate detachment. However, once the initial adhesion, indicated by the first large spike in force, has been overcome, membranes detach soon after due to the lack of additional sites of attachment.

**Figure 4.**
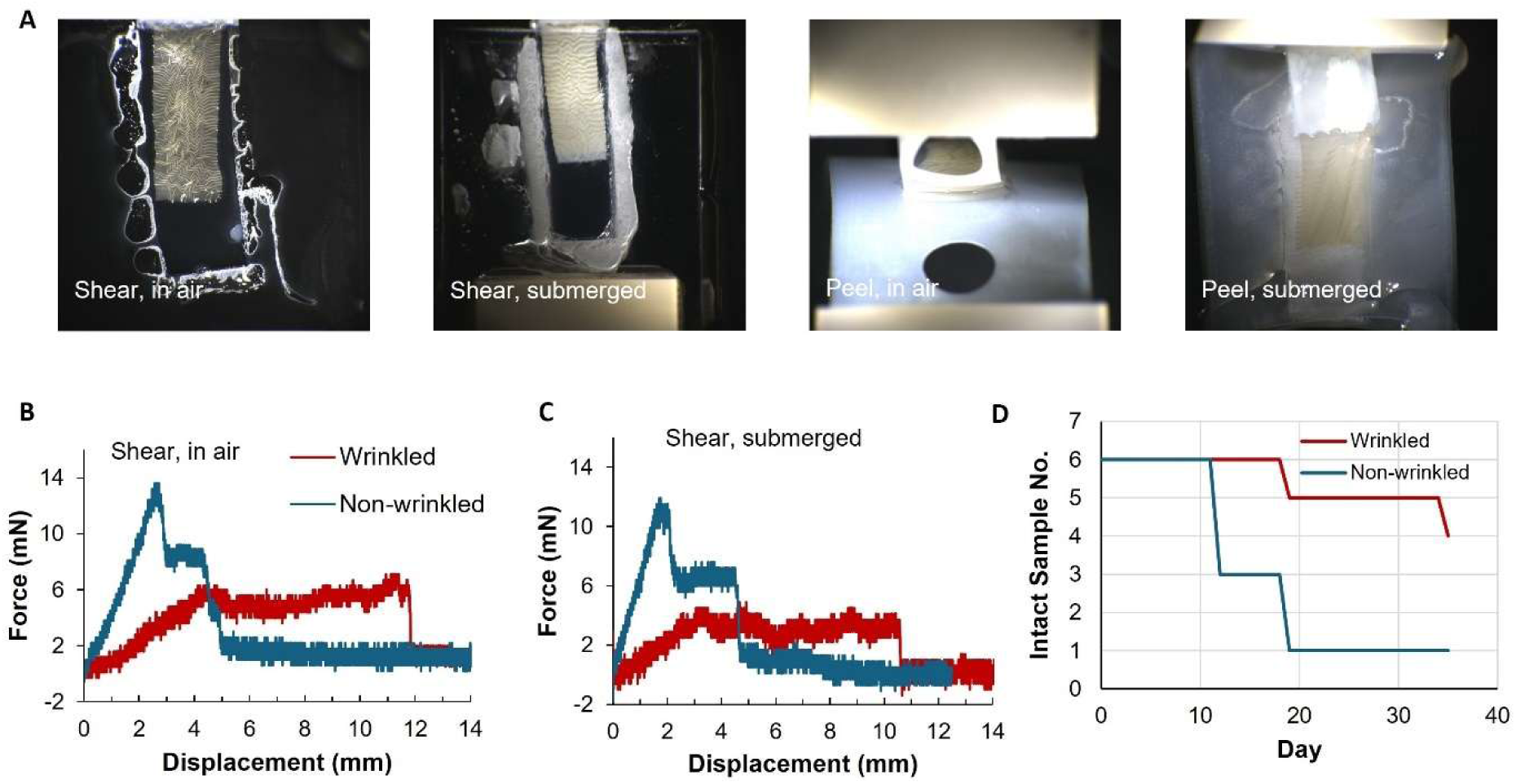
Influence of wrinkle formation on interfacial adhesion. (**A**) Peel and shear tests were conducted on wrinkled and non-wrinkled bilayer samples (sample dimensions: 20 mm x 6.5 mm x 1 mm) either submerged in PBS or in air. (**B-C**) Representative force curves of wrinkled and non-wrinkled samples obtained for shear tests conducted in air (B) and in PBS (C). (**D**) Number of intact bilayer samples exposed to mild agitation on a rocking platform (150 rpm) in a submerged state at 37 °C.

In peel tests, both wrinkled and non-wrinkled samples demonstrated similar force curves both in air and in a submerged state (Supporting Information, **Fig. S3**). This is an expected outcome since peeling force is directly related to the width of the adhesion interface, rather than the area. Consequently, measured values were lower than what was observed for shear tests, especially for submerged samples, with no major spikes in force during detachment. The peel test results thus imply that the presence of interfacial wrinkles does not substantially add to bilayer adhesion at individual local contact zones (e.g. the peeling front). Fluid transportation from the hydrogel surface to the dry membrane could provide the primary interfacial interactions [40]. However, an assessment of the long-term stability of adhered bilayers provided by wrinkle formation showed that wrinkled cfGel-Membranes remained attached over a longer period of time compared to their non-wrinkled counterparts when exposed to mild agitation on a rocking platform (150 rpm) in a submerged state at 37 °C (**Fig. 4D** and Supporting Information **Fig. S4**), thus underscoring the important role of wrinkles in maintaining structural integrity. This is particularly relevant for our application of the bilayer as a model for skin tissue engineering, as the wrinkled surface needs to remain stable over a prolonged culture period without detachment.

### DEJ-inclusive scaffolds as co-culture platforms for HEKs and HDFs

To verify the potential of our bilayer system as a platform for skin tissue engineering, we first evaluated the suitability of cfGel-Membranes to act as a growth substrate for HEKs. Metabolic activity of HEKs was measured over 7 days of culture to indirectly assess cell proliferation (**Fig. 5A**). HEKs demonstrated increased metabolic activity on day 4 and 7 when compared to the initial value obtained on day 1 for passage 5 cells, though results varied with increasing passage number (Supporting Information, **Fig. S5**). The results suggest that using D-fructose to crosslink gelatin membranes via a Maillard reaction-based process constitutes a viable option for the fabrication of nanofibrous growth substrates for the culture of keratinocytes. We established HDF-HEK co-cultures using bilayer scaffolds as illustrated in **Fig. 1C**. After a submerged incubation time of 1-3 days, scaffolds were lifted to the air interface and cultured for further 3-4 weeks. Histological evaluation of models after co-culture revealed a clear separation of the epidermal and dermal layers by nanofibrous membranes without visible signs of cell infiltration into the other compartment, thus restricting HDFs and HEKs to their respective cutaneous layers (**Fig. 5B**). Visualization of membranes through incorporation of fluorescently labeled dextran revealed retention of wrinkles throughout the entire culture period (**Fig. 5C**). The wrinkles further demonstrated persistent tight contact with the underlying hydrogel, which is in agreement with the data obtained from adhesion tests. The result implies that nanofibrous membranes can support the formation of an epidermal layer by enabling seeding of HEKs at any given point without cells falling through or penetrating into the scaffold, thus not requiring previous deposition of a dermal ECM by HDFs. H&E stainings show the formation of epithelia of variable thickness and stratification depending on their location (**Fig. 5B** and Supporting Information **Fig. S6**). Epithelial layers were especially thin at the peaks of formed wrinkles, consisting mostly of a cornified layer of flattened keratinocytes, whereas cells located within the pockets formed during wrinkling demonstrated a more rounded morphology. This phenomenon was observed for both bilayer models with cfGel-Hydrogels of 1 mm and 0.2-0.3 mm thickness. It is therefore unlikely that insufficient diffusion of nutrients towards basal keratinocytes during air-lift culture constituted a decisive factor. Due to the rapid initiation of wrinkling upon contact with cell culture media during HEK seeding, cells are not afforded sufficient time to adhere to membranes, resulting in most of them accumulating in valleys with only a small number remaining on the peaks and slopes of the formed wrinkles. This likely interfered with the formation of a confluent HEK monolayer prior to air-lift culture and might have affected epithelial maturation. Instead of organizing into a homogeneous monolayer that eventually matures into a multi-layered stratified epidermis, HEKs primarily grew out of pockets and quickly underwent terminal differentiation into a cornified layer. This is further corroborated by immunohistological stainings that showed the expression of cytokeratin 10 as a marker of the stratum corneum in cases where HEKs covered a larger area (**Fig. 5C**). Although laminin, as a marker of the basement membrane typically expressed by basal keratinocytes, does appear to be correctly localized on the epidermal side of the nanofibrous membrane in several instances, it could not always be detected despite the presence of HEKs. Preliminary data using an improved experimental setup that allows for keratinocytes to adhere onto membranes and form confluent monolayers prior to wrinkling exhibits notably thicker epithelia and clear differentiation of the cornified layer with the expression of layer-specific markers such as loricrin (**Fig. 6** and Supporting Information, **Fig. S6**). Furthermore, H&E staining indicated a higher density of keratinocytes stained positively for nuclei situated within epidermal pockets, implying a potential influence of microtopographical features on keratinocyte differentiation.

**Figure 5.**
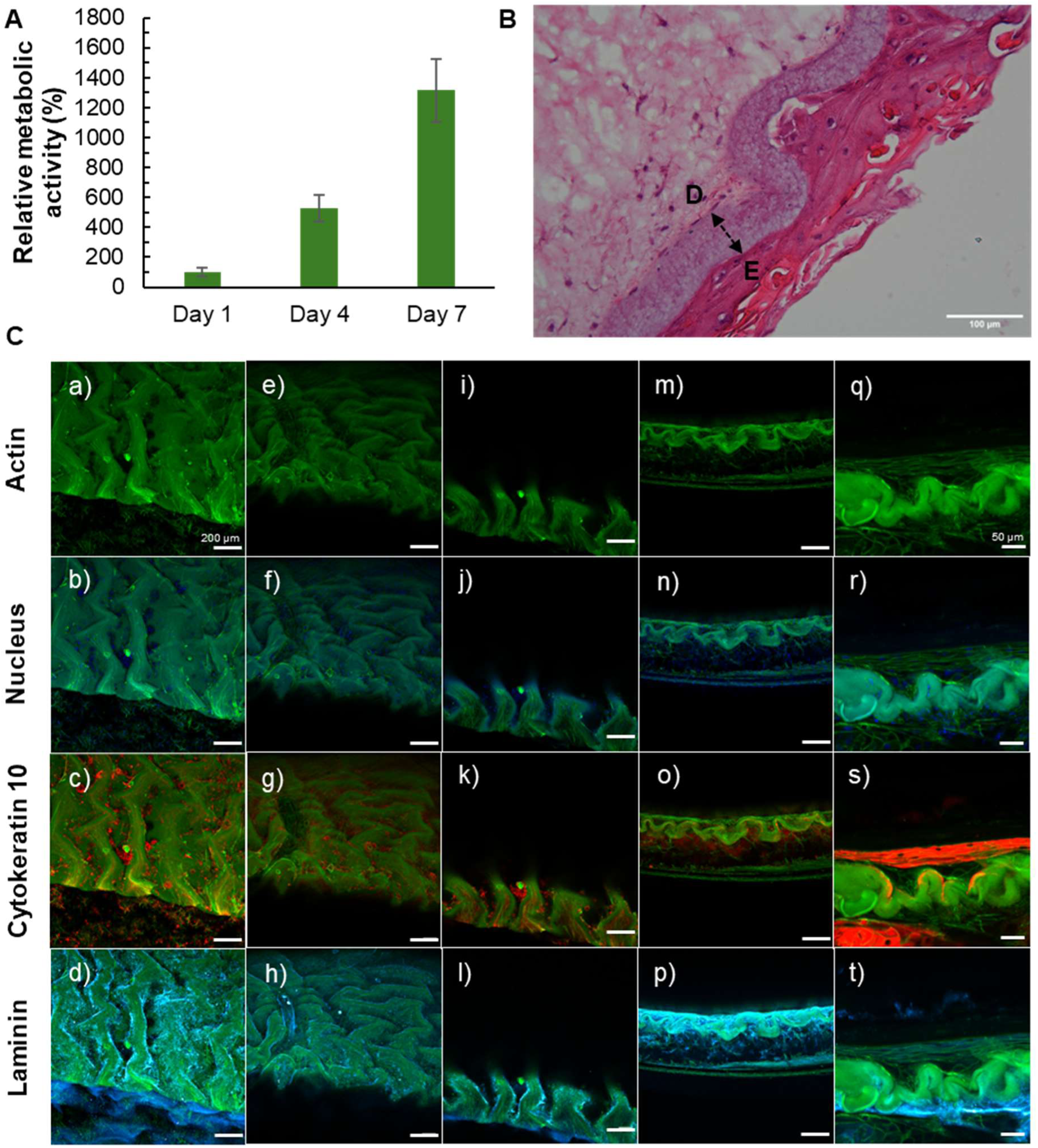
Co-culture and tissue evaluation. (**A**) Results of MTT assay reveal an increase in metabolic activity of HEKs (passage #5) cultured on nanofibrous membranes over time. Absorbance (550 nm) was normalized to values obtained on day 1 (100%). Error bars indicate SD. *n = 3*. (**B**) H&E staining of bilayer model shows clear separation of epidermal (E) and dermal (D) layers by nanofibrous membranes (arrow). No visible penetration of cells into the fibrous matrix was detected. (**C**) Immunological staining using laminin and cytokeratin 10 as markers of the basement membrane and stratum corneum respectively. Green fluorescence of the membrane is due to the inclusion of fluorescently labeled dextran. Images a-h show the surface of the models at different angles, whereas i-t show cross-sections. a-p: Scale bar = 200 µm, q-t: Scale bar = 50 µm.

**Figure 6.**
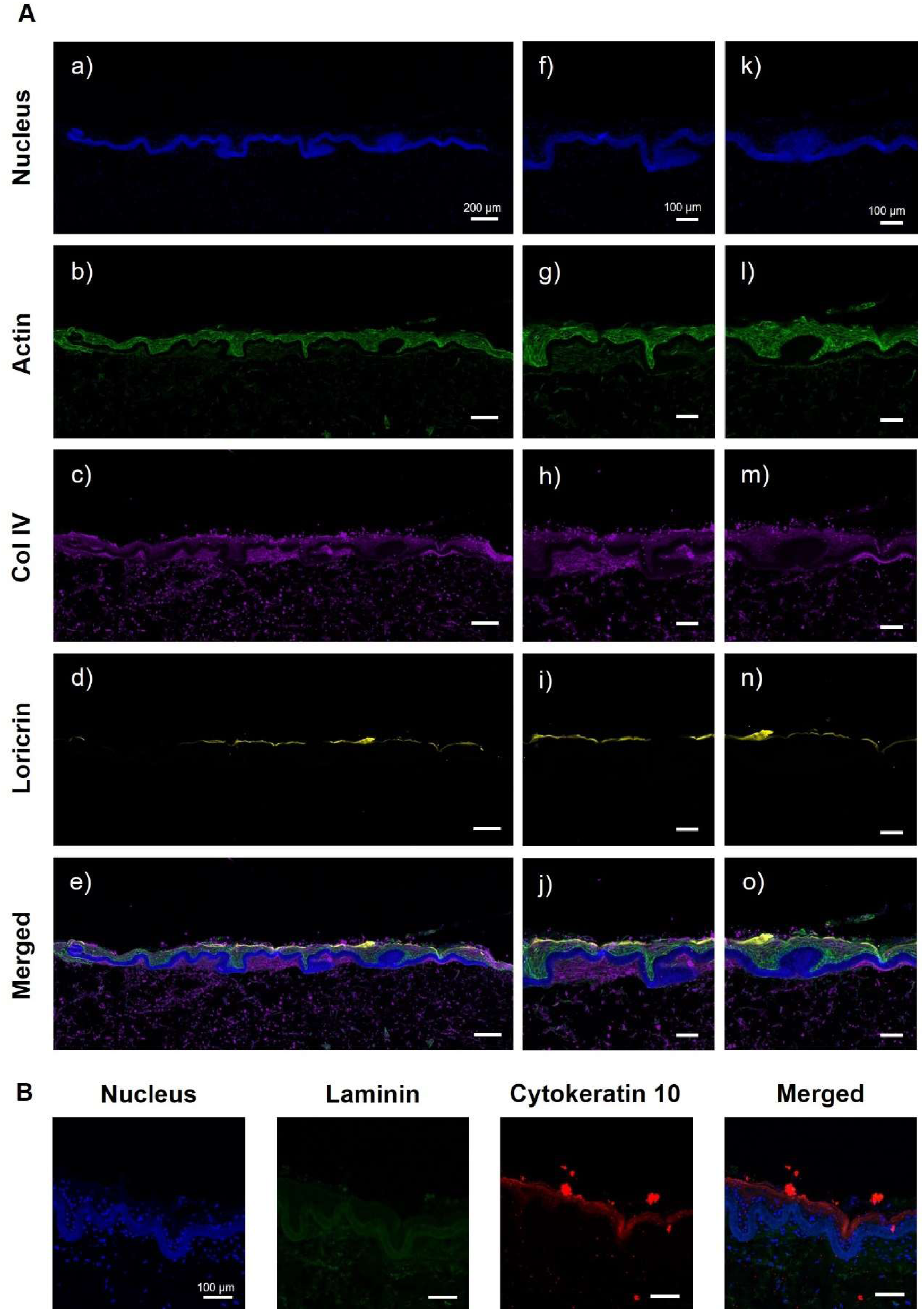
Improved epithelial maturation. (**A**) Allowing HEKs to form confluent monolayers prior to wrinkling results in the formation of thicker epithelial layers. Expression of loricrin and cytokeratin 10 (**B**) as markers of the cornified layer indicates improved epidermal maturation. Expression of the basement membrane marker collagen IV on the other hand appears to be less specific, with areas of higher intensity being located both on the dermal and epidermal side of cfGel-Membranes. DAPI staining was used to visualize cfGel-Membranes. Images show cryosections of bilayer models at full-length (A: a-e) and zoomed-in sections (A: f-o, B). A: a-e: Scale bar = 200 µm, A: f-o and B: Scale bar = 100 µm.

### Conclusion and outlook

We developed a bilayer scaffold composed of a nanofibrous membrane and an underlying hydrogel support that exhibits DEJ-like morphological features for the purpose of *in vitro* skin tissue fabrication. To mimic the undulated topography of the DEJ, we exploited the controllable swelling behavior of cfGel-Hydrogels to facilitate volumetric changes of the scaffold’s dermal component, resulting in buckling instabilities between the two layers due to geometrical constraints imposed by the nanofibrous membrane. Increasing mechanical stress eventually cumulates in the formation of surface wrinkles that demonstrate topographical similarity to the rete ridges and dermal papillae of the DEJ. This method represents a facile way to dynamically introduce microtopographical features in the presence of cells, allowing them to be exposed to mechanical stimuli that would otherwise not be present in pre-patterned static surfaces. In this context, the functional relevance of the DEJ-inspired interface is assessed in terms of interfacial stability, spatial regulation of tissue architecture, and support of epidermal stratification and maturation. Future research directions may include the evaluation of additional functional aspects of the skin models, such as barrier and transport performance.

Analysis of surface roughness parameters as well as size and numbers of wrinkles suggest a high degree of consistency across different samples. It is noteworthy that the random formation of wrinkles on a given sample can result in notable differences in wrinkle number and dimensions observed for cross-sections depending on the position of the cut. Therefore, additional analyses should be conducted in the future to further elucidate the influence of swelling time on the generation of topographical features. In addition, computational simulations, which have been employed in the past to predict the formation of surface wrinkles and explain the development of rete ridges, could enable a more precise manipulation of system parameters to obtain specific surface patterns in the future [30, 56]. Overall, although the method is currently limited in its ability to precisely reproduce specific surface topographies, the simplicity of its application and the compatibility of the swelling/deswelling-based approach with a broad range of 3D-shapes and sizes of samples constitutes an advantage to alternative patterning methods relying on mechanical stretch or advanced manufacturing techniques. Optimization of process parameters such as swelling/deswelling conditions or layer thickness, could result in the formation of wrinkles with specific dimensions that mimic the rete ridges at different body locations, ages, or pathological conditions. Continued investigation will provide deeper insights into the long-term stability of the bilayer scaffolds and the functional performance of the engineered skin models.

## Acknowledgements

T. H. acknowledges Julia Hau for her assistance with histology. Some images were partially created using Biorender (Fig. 1c).

## Funding

This work was supported by Empa’s Directorate Board (SKINTEGRITY.CH collaborative research program).

## Conflict of interest

Tobias Hammer reports financial support was provided by Skintegrity.CH. The authors are named inventors on patent application WO2026037844A1. The authors declare no other competing financial interest.

## Materials and Methods

### Hydrogel preparation

Hydrogels were prepared by exposing precursor solutions containing equal amounts of norbornene-functionalized cold-water fish gelatin (cfGel-NB) and thiol-functionalized cold-water fish gelatin (cfGel-SH) at a final concentration of 5% (w/w) and 0.05% (w/v) Irgacure 2959 to 365 nm UV-irradiation for 2 min, as described in a previous report [32]. Briefly, cfGel-Hydrogel precursor solutions were prepared by dissolving lyophilized cfGel-NB and cfGel-SH in 1x PBS at room temperature, followed by addition of 2-Hydroxy-4′-(2-hydroxyethoxy)-2-methylpropiophenone (Irgacure 2959, stock solution 0.5% (w/v) in ultrapure water). Precursor solutions were then cast into various PDMS/glass-based molds depending on the respective application and subsequently exposed to UV-light at 365 nm for 2 min using a 12W (VL-206.BL, 2×6W, 365 nm) UV light source. Lubrication of the molds with water or PBS allowed for better removal of its components without damaging the gels. To remove hydrogels from molds with a glass support, we found a cell-scraper with a rubber blade to be the most suitable option.

### Evaluation of swelling behavior

Swelling ratio (SR) of hydrogels after swelling-deswelling in 0.25 M sucrose solution and 1x PBS respectively was evaluated by recording both changes in weight and diameter at specified timepoints (t). Hydrogels were prepared using a custom-made mold consisting of a rectangular sheet of PDMS of 1 mm thickness containing holes of 6 mm diameter pressed onto a glass slide. Prior to UV-irradiation, the mold was topped with a second sheet of PDMS to ensure the resultant hydrogels would exhibit a flat surface on both sides. After recording their initial weight and diameter using a KERN precision balance and an Aerospace 150 mm Digital Vernier Caliper respectively, hydrogels were submerged in 3 mL of sucrose solution for 15 min, 30 min, 45 min or 60 min. Swollen hydrogels were once again measured for both weight and diameter before being transferred into 3 mL of PBS. During deswelling, weight and diameter were periodically measured after 30 min, 1 h, 2 h, 4 h, 6 h and 24 h. Throughout both incubation periods, hydrogels were kept in 12-well TCC plates at 37 °C. To prevent excessive evaporation of liquid, well plates were sealed with parafilm overnight. After measuring the final weight and diameter, the swelling ratio (SR) of the hydrogels was determined using the following formula:

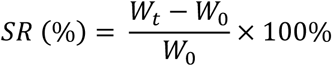

with *W_0_* representing the initial weight of as-prepared hydrogels and *W_t_* representing the measured weight at timepoint *t*.

### Preparation of nanofibrous cfGel-Membranes from cold-water fish gelatin

Nanofibrous membranes were fabricated by electrospinning and crosslinking unmodified cold-water fish gelatin with reducing sugars according to previous descriptions found in literature [33, 34]. The spinning dope was prepared by dissolving 4 g of cold-water fish gelatin and 400 mg of D-fructose in 10 mL ultrapure water at room temperature and either used directly afterwards or stored at 4 °C until further use. The dope solution was subsequently loaded into 1 mL or 2 mL syringes (Braun, Injekt® Luer Solo) fitted with a 21G blunt tip needle (Braun, Sterican®) and mounted onto a programmable single channel syringe pump (WPI, Model No. Aladdin-1000) spaced at a distance of 15 cm between the needle tip and the collector surface. The spinning was conducted at a fixed flow rate of 0.4 mL/h and an applied voltage of 15 kV at 25 °C and ∼46% relative humidity in a controlled environment. Fibers were collected onto a sheet of aluminum foil mounted on a rolling stainless drum (13.5 cm length, 5 cm diameter) rotating at 60-90 rpm. After extruding a total volume of 1 mL, fiber mats were collected and stored in a desiccator to allow for complete drying prior to being incubated in an oven at 140 °C for 8h to facilitate crosslinking. Finally, crosslinked fiber mats were stored in a desiccator at room temperature until further use. For observation of nanofibrous membranes during confocal microscopy, ∼1-2 mg/mL of fluorescently labeled dextran (FD70, Sigma-Aldrich) was added to the spinning dope. Average fiber diameters were calculated from a total of 736 individual fibers measured in ImageJ across *n = 3* samples.

### Formation and characterization of surface wrinkles

To evaluate the effect of swelling time on the morphology of resultant surface wrinkles, disc-shaped hydrogels with a diameter of 6 mm and a thickness of 1 mm were immersed in 3 mL of sucrose solution for 0 min, 15 min, 30 min, 45 min or 60 min to induce varying degrees of volumetric expansion. Swollen hydrogels were subsequently recovered and blotted free from excessive surface liquid using a cotton swab (Copan). Removing excess liquid establishes better contact between the hydrogel and the electrospun membrane and prevents the membrane from being preemptively hydrated before getting in contact with the gel. Circular membranes with 6 mm diameter (for 0 min and 15 min gels) and 8 mm diameter (for 30 min, 45 min and 60 min gels) were punched out using biopsy punches (KAI Medical) and carefully separated from the aluminum foil support using tweezers before being placed on top of hydrogels using a PDMS stamp. Membrane-covered hydrogels were subsequently transferred into 3 mL of PBS to de-swell overnight.

Characterization of the surface topography of wrinkled samples was conducted using a KEYENCE VHX-7000 optical microscope. Before analyzing the surface, excessive liquid was removed by contacting the samples laterally with a cotton swab without disturbing or altering topographical features. Surface profiles of samples were obtained by imaging the samples from a top shot perspective. Evaluation of the surface roughness parameters *Sa, Sz, Sq, Ssk, Sku, Sp* and *Sv* was performed using a Gaussian filter at a cut-off frequency of λc = 8 mm, 2.5 mm, 1 mm, 0.8 mm and 0.5 mm.

Wrinkle valleys were measured in ImageJ using the sample profiles obtained from Keyence analysis. The number of wrinkles measured for each sample were 48 for 0 min, 61 for 15 min, 71 for 30 min, 70 for 45 min, and 76 for 60 min. For each pre-swelling time, six samples were analyzed (one profile per sample). The number of wrinkles for each pre-swelling condition was determined by averaging the wrinkle number per millimeter for each sample. Statistical analysis of wrinkle depths and numbers was conducted using a Kruskal-Wallis and Mann-Whitney U test and Student’s t-test respectively.

### Evaluation of membrane adhesion with peel -and shear tests

To investigate the influence of surface wrinkles on the adhesive strength between electrospun membranes and cfGel-Hydrogels, both peel -and shear tests were conducted on wrinkled and non-wrinkled samples, both in air and submerged in 1x PBS at room temperature. Surface wrinkles were introduced by swelling rectangular-shaped cfGel-Hydrogels (20 mm x 6.5 mm x 1 mm) in 3 mL of ultrapure water in 6-well TCCP plates for 1 h at room temperature, followed by membrane attachment and subsequent deswelling in 1x PBS for at least 3 h. Electrospun membranes were prepared by cutting out rectangular shapes of 39 mm x 6.5 mm size using a custom-made setup consisting of two razor blades fixed at a distance of 6.5 mm. The contact area between membranes and hydrogels for both swollen and non-swollen hydrogels was approximately 20 mm x 6.5 mm. The remaining overhang of the membranes was fixed between two pieces of sticky tape and a hole with a diameter of 5 mm was punched into the far end using a biopsy puncher.

Both peeling and shearing tests were conducted using a BioTester biaxial test machine (CellScale) at a constant rate of 0.25 mm/s. For shear tests, hydrogels were adhered onto glass supports using instant glue and fixed to the gooseneck using the clamp mounting system. The membrane was fixed onto the opposing gooseneck by threading the pin of the calibration spring bracket through the hole in the sticky tape-fixed part. For shear tests performed on samples submerged in PBS, hydrogels were instead adhered to the bottom side of the glass supports to allow for complete submersion within the fluid chamber without it being contacted by any elements of the mounting system, which could interfere with the measurement.

Peeling tests were conducted in a T-peel test-like manner by adhering hydrogels onto a 2-layered piece of sticky tape using instant glue and fixing both the sticky tape support as well as the sticky tape-reinforced membrane overhang into the clamp mounting system.

### Deswelling force measurement

To measure the amount of force exerted by cfGel-Hydrogels during deswelling, rectangular hydrogels (30 mm x 6.5 mm x 1 mm) were first swollen in 3 mL of ultrapure water for 1 h at room temperature. Afterwards a biopsy puncher was used to introduce two holes of 5 mm diameter into each end of the hydrogels, making sure to leave enough material between the holes and the borders to prevent breaking of the hydrogels during deswelling. Hydrogels were subsequently mounted onto a BioTester biaxial test machine (CellScale) by threading the pins of the calibration spring brackets through the holes of the swollen hydrogels and lowering the fixed samples into a fluid chamber filled with 1 x PBS. To prevent the influence of prior mechanical stretch on the measured force, the distance between the two pins was set to be shorter than the length of the swollen hydrogel. The measurement was conducted for 1 h with a data measurement frequency of 1 Hz and an image capture rate of 0.01 Hz.

### HEK proliferation assay

HEK proliferation was evaluated by measuring metabolic activity over time via MTT assay. Nanofibrous membranes with a diameter of 10 mm were placed into the wells of a 48-well plate using a PDMS stamp. Prior to membrane placement, PBS was added and subsequently removed from the wells to slightly wet the bottom surface and cause dry membranes to stick upon contact. This allowed membranes to remain adhered to the bottom of the wells without further fixation. Afterwards, membranes were covered with Prime Epithelial Proliferation Medium (Cnt-Pr, CELLnTEC) prior to cell seeding. Primary human epidermal keratinocytes (HPEKp, CELLnTEC, Lot. EI2009089) at passage #5, #6, and #7 were seeded onto nanofibrous membranes at a density of 40’000 cells/well and cultured for 1 week with media changes every other day. At day 1, 4, and 7, cells were incubated with 200 µL of Cnt-Pr medium supplemented with 50 µL of MTT solution (stock: 5 mg/mL in PBS) for 4 h before MTT formazan was extracted with 100 µL of extraction solution (90% ethanol + 10% HEPES-NaCl) and measured at 550 nm using a Mithras LB 943 Multimode Microplate Reader.

### Cell culture

Primary human dermal fibroblasts (HDFs, CELLnTEC, Lot. EB1104281, pooled juvenile donors) were cultured in Dulbecco’s Modified Eagle’s Medium (DMEM, Sigma-Aldrich, D5796) supplemented with 10% (v/v) fetal bovine serum (Sigma-Aldrich, F9665, Lot. 0001655440), 1% penicillin-streptomycin (Sigma-Aldrich, P4458-100ML, Lot. 0000192944), and 1% L-glutamine (Sigma-Aldrich, G7513-100ML, Lot. RNBM0086) in a humidified environment at 5% CO_2_ and 37 °C. HDFs were used up to passage #10. Cells were cultured in standard tissue culture plastic flasks (TPP, Product No. 90076) with media changes being conducted twice a week until 80%-100% confluency. Following trypsinization, cells were collected in cell culture medium, pelleted and reconstituted in cfGel-Hydrogel precursor solution at a density of 2’000’000 cells/mL for cell encapsulation purposes. Cell-laden hydrogels were molded using a custom-made mold consisting of a rectangular sheet of PDMS of either 1 mm or 0.2-0.3 mm thickness containing holes of 8 mm or 12 mm diameter pressed onto a glass slide. After casting the precursor solution into each well, the loaded mold was covered with a second sheet of PDMS prior to crosslinking. Hydrogels were recovered from the glass support with a rubber cell scraper and placed in 12-well plates with 2 mL of culture media. Encapsulated HDFs were typically cultured for 1 week before bilayer assembly and seeding of keratinocytes but have also been used as early as the day after encapsulation.

Primary human epidermal keratinocytes were cultured in Cnt-Pr medium in a humidified environment at 5% CO_2_ and 37 °C up to passage #7. Cells were cultured in standard tissue culture plastic flasks (TPP, Product No. 90076) with media changes being conducted 3x a week until ∼60-70% confluency.

Before seeding of primary human epidermal keratinocytes, HDF-inclusive cfGel-Hydrogels were swelled in 0.25 M sucrose solution for 1 h at 37 °C followed by attachment of nanofibrous membranes with diameters of 8 mm using a PDMS stamp and placement of cloning cylinders around membranes. HEKs were detached from flasks using accutase, pelleted, and reconstituted in CnT-Prime Full Thickness 3D Airlift Medium (CnT-PR-FTAL5, CELLnTEC) before being seeded directly onto swollen scaffolds at a density of 300’000 cells/sample. Scaffolds were incubated at 37 °C for 30-60 min to allow for cells to adhere prior to wells being carefully supplemented with 1-2 mL of CnT-PR-FTAL5 to slightly cover the top of the bilayer scaffolds. After submerged culture for 1-3 days, samples were lifted to air-lift culture by placing scaffolds into transwell inserts and placing them into 12-well plates fitted with a CnT Spacer Plate (CnT-SP, CELLnTEC) containing 1.8 mL of CnT-PR-FTAL5. Scaffolds were cultured for 3-4 weeks with media being changed 3x per week.

### Histology

For the preparation of histological sections, bilayer scaffolds were first fixed o/n in 5% neutral buffered formalin (Sigma-Aldrich, HT501128) at 4 °C, followed by another o/n incubation period in a 30% (w/v) sucrose solution at 4 °C. Cryomolds (Tissue-Tek, Sakura) were used to embed scaffolds in optimal cutting temperature (OCT, Tissue-Tek, Sakura) compound before freezing in 100% ethanol cooled down with a liquid nitrogen/dry ice slurry. Samples were then transferred and stored at −80 °C until further use. Samples were cut into 10 μm thick sections using a Dakewe CT520 cryostat microtome and mounted onto Superforst Plus Adhesion Microscope Slides (Epredia, J1800AMNZ). Sections were stained with hematoxylin (Harris HTX, Biosystems, 41-1011-00) and eosin (Eosin Y Solution, Alcoholic, Sigma-Aldrich, HT110132-1L) using a Slide Stainer MYREVA SS-30. After drying, sections were mounted using ROTI®Histokitt (Roth, Art. No. 6638.1). Alternatively, sections were stained for immunofluorescence analysis.

### Immunofluorescence staining and analysis

Immunofluorescence analysis was conducted by fixing bilayer scaffolds in 5% neutral buffered formalin o/n at 4 °C before cutting samples in half and permeabilizing them with 0.1% Triton X-100 (Sigma-Aldrich, T8787-100ML) for ∼15 min. Samples were subsequently washed 3x with DPBS for 1 min before either being stored at 4 °C or directly processed further. Blocking was performed in 3% bovine serum albumin (BSA in DPBS, Sigma-Aldrich, A9647-50G) for at least 1h at room temperature or o/n at 4 °C before samples were incubated in primary antibody solution o/n at 4 °C. After secondary antibody incubation for 1h at room temperature in the dark, samples were washed 3x with DPBS for 1 min. All antibodies were diluted in DPBS containing 1% fetal bovine serum (dilution factor indicated below), with DAPI (D9542, Sigma-Aldrich) being diluted by 1:1000.

#### Primary antibody

- Anti-Cytokeratin 10 antibody [EP1607IHCY]
- Cytoskeleton Marker; Abcam; 1:200
- Anti-Laminin beta 1 antibody [LT3]; Abcam; 1:200
- Anti-Loricrin antibody [EPR7148(2)(B)]; Abcam; 1:200
- Mouse anti-Collagen IV, clone MC4-HA (Monoclonal); Monosan; 1:20

#### Secondary antibody

- Goat anti-Rabbit IgG (H+L) Secondary Antibody, Alexa Fluor 555; Invitrogen; 1:200
- Goat IgG anti-Rat IgG (H+L)-Cy5; Jackson; 1:200
- Goat anti-Rabbit IgG (H+L) Highly Cross-Adsorbed Secondary Antibody, Alexa Fluor 633; Invitrogen; 1:200
- Goat anti-Rabbit IgG H&L (Alexa Fluor 488); Abcam; 1:200
- Goat anti-Mouse IgG (H+L) Cross-Adsorbed Secondary Antibody (Alexa Fluor 555); Invitrogen; 1:200

#### Conjugated antibody

- Phalloidin F-actin; Alexa Fluor 488; Invitrogen; 1:200
- Phalloidin F-actin; Alexa Fluor 633; Invitrogen; 1:200

#### Data availability

Data will be made available on request.

## Notes

### Summary of Updates

The last author's middle name was corrected.

