## supporting information for "A dermal-epidermal junction-inclusive skin model enabled by controllable hydrogel swelling"

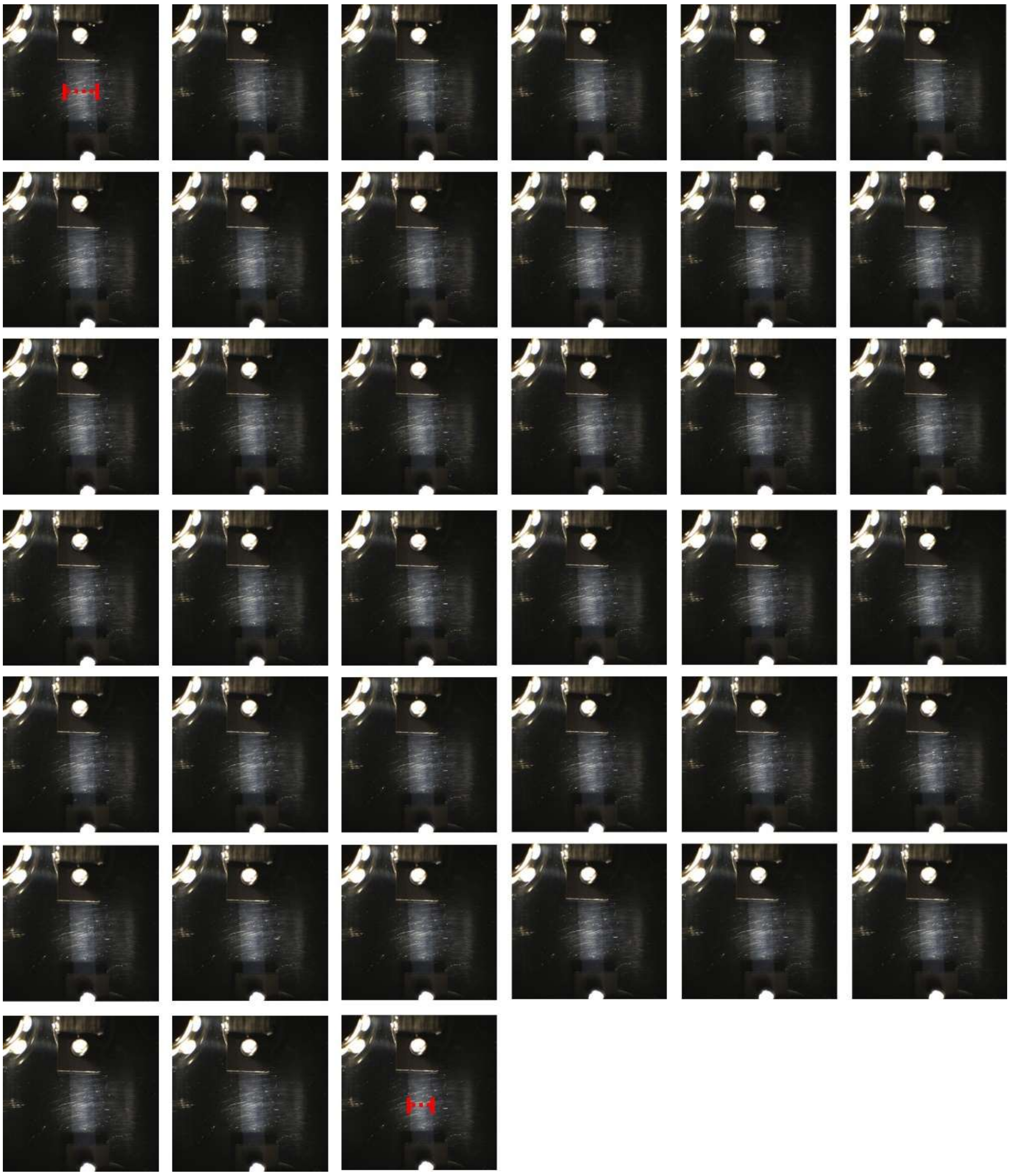

**Figure S1. Experimental setup to measure force generated by deswelling cfGel-Hydrogels over time.** Top left to bottom right: Visualization of the deswelling process of cfGel-Hydrogels submerged in PBS. Images were taken every 100s.

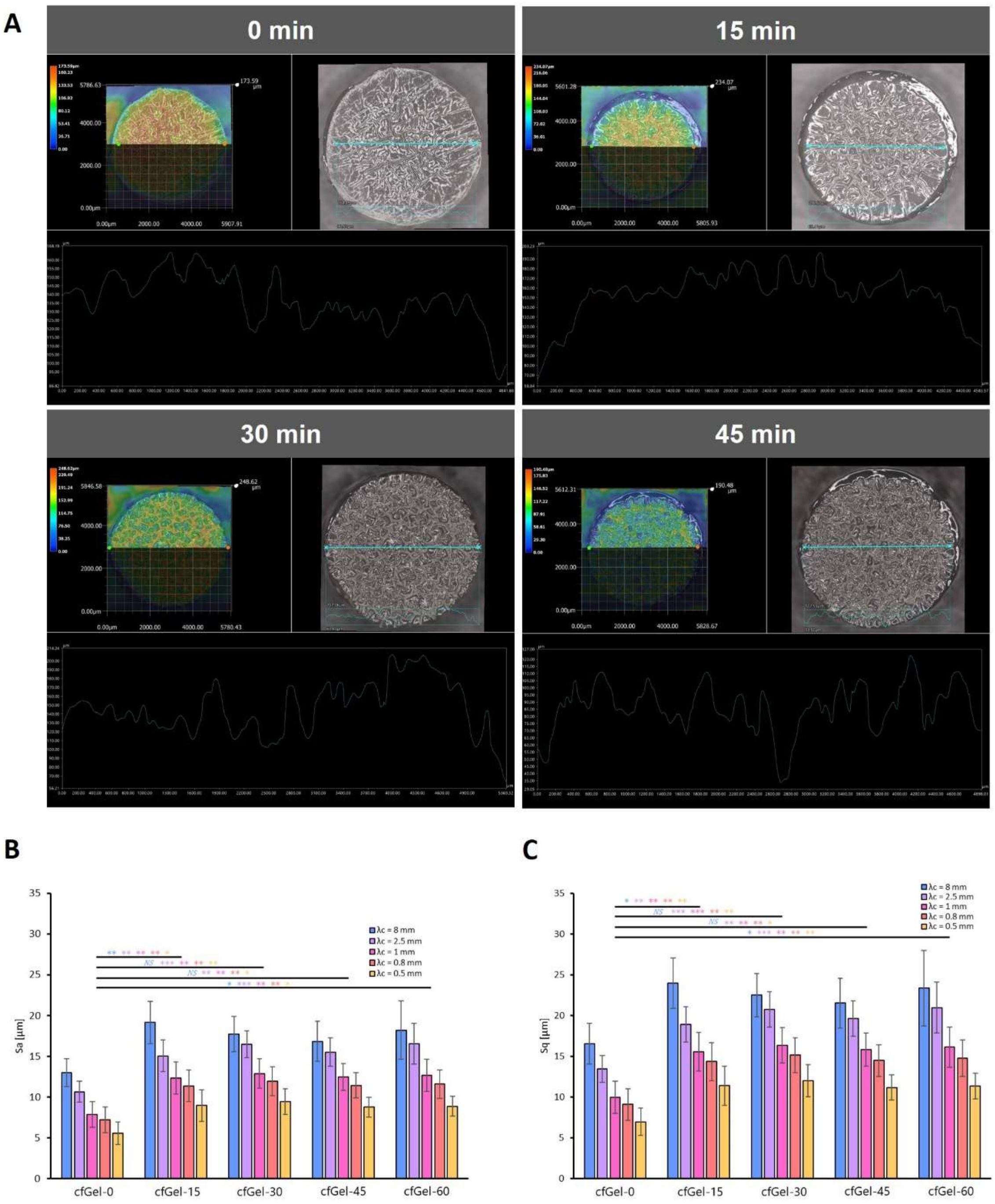
A
0 min
15 min
30 min
45 min
B
Sa [µm]
λc = 8 mm
λc = 2.5 mm
λc = 1 mm
λc = 0.8 mm
λc = 0.5 mm
NS
cfGel-0
cfGel-15
cfGel-30
cfGel-45
cfGel-60
C
Sq [µm]
λc = 8 mm
λc = 2.5 mm
λc = 1 mm
λc = 0.8 mm
λc = 0.5 mm
NS
cfGel-0
cfGel-15
cfGel-30
cfGel-45
cfGel-60

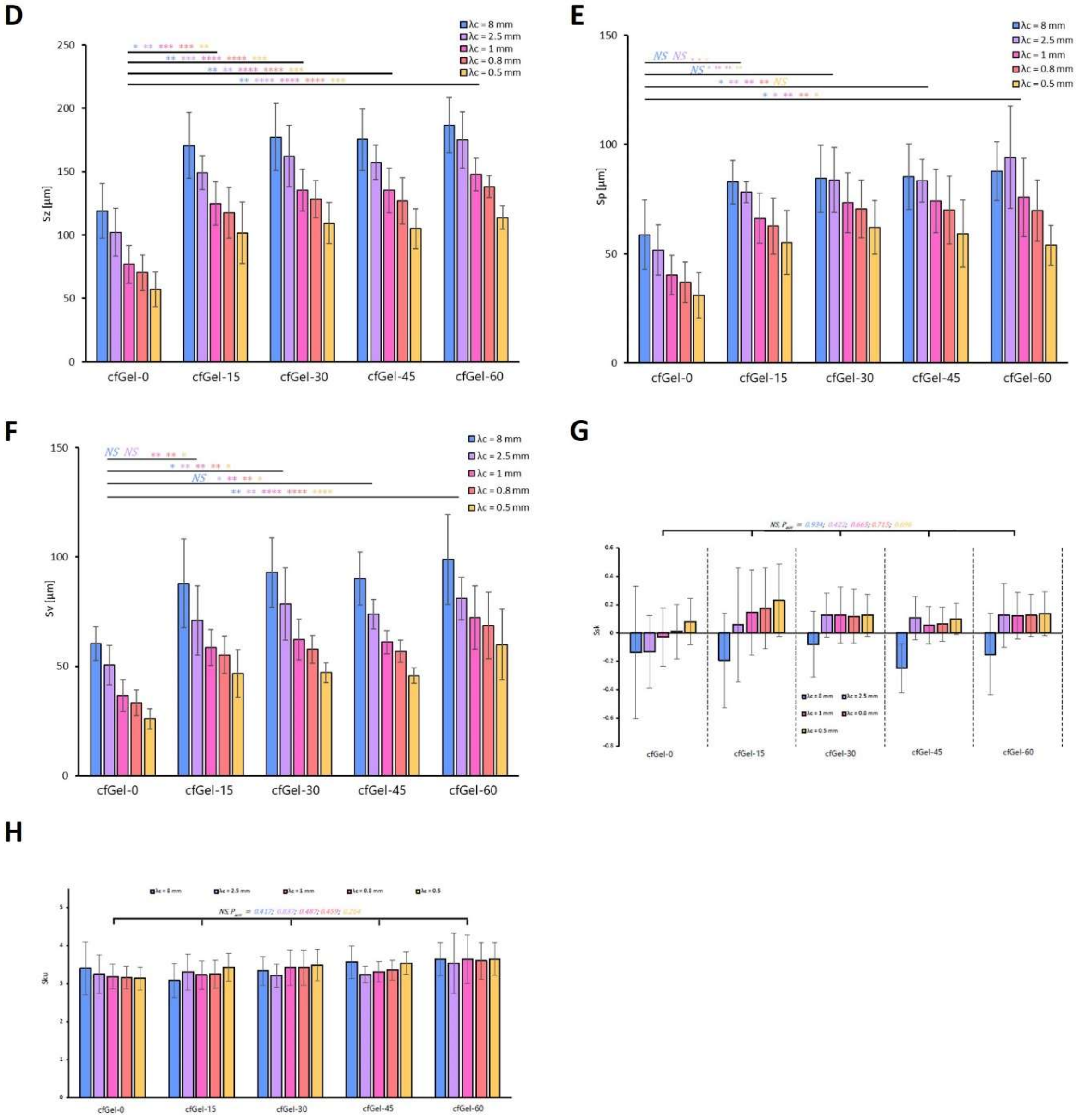


**Figure S2. Surface topography and profile analysis of bilayer scaffolds.** (**A**) Results of surface topographical analysis of scaffolds with pre-swelling times of 0 min, 15 min, 30 min, and 45 min. Arithmetic mean height *Sa* (**B**), root mean square height *Sq* (**C**), maximum height *Sz* (**D**), maximum peak height *Sp* (**E**), maximum pit depth *Sv* (**F**), skewness *Ssk* (**G**), and kurtosis *Sku* (**H**) of samples with different pre-swelling times under application of a Gaussian filter with different cut-off values. Error bars depict SD, $n = 6$. The p-values were obtained by Tukey HSD test, following one-way ANOVA. Only data of identical cut-off were compared (colour-coded). *P < 0.05, **P < 0.01, ***P < 0.001, ****P < 0.0001, NS = not significant.

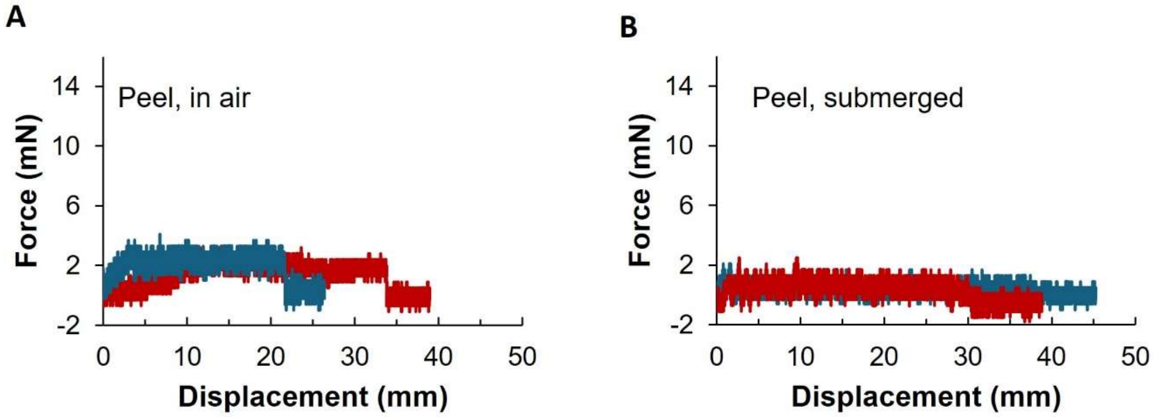


**Figure. S3** Representative force curves of wrinkled and non-wrinkled samples obtained for peel tests conducted in air (**A**) and in PBS (**B**).

| Date | Wrinkled | Non-wrinkled |
|---|---|---|
| 28.04.2025 | 6 attached | 6 attached |
| 01.05.2025 | 6 attached | 6 attached |
| 12.05.2025 | 6 attached | 3 attached<br>3 detached |
| 19.05.2025 | 5 attached<br>1 partially detached | 1 attached<br>5 detached |
| 26.05.2025 | 5 attached<br>1 partially detached | 1 attached<br>5 detached |
| 02.06.2025 | 4 attached<br>2 detached | 1 attached<br>5 detached |
| 10.06.2025 | 2 attached<br>4 detached | 1 attached<br>5 detached |

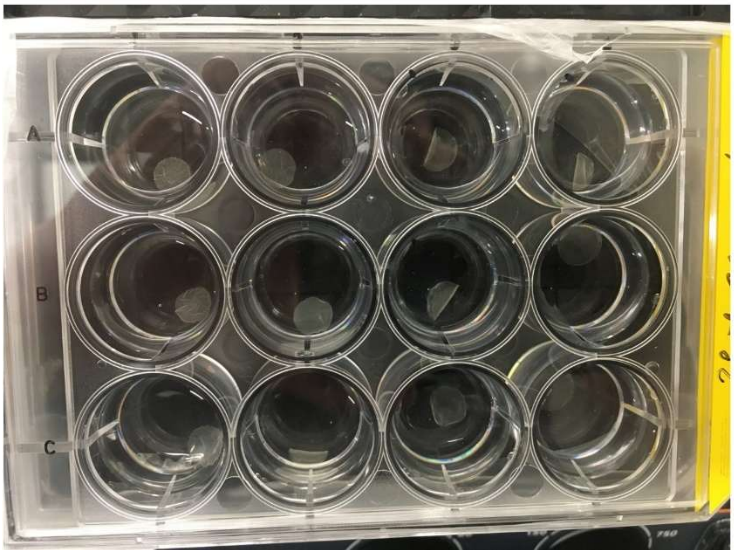


**Figure S4. Evaluation of long-term stability of wrinkled and non-wrinkled bilayer scaffolds.** Scaffolds were submerged in PBS and incubated at 37°C on a rocking platform at 150 rpm. PBS was changed every week. Wrinkled bilayers exhibited increased stability, with detachment of membranes occurring at later times and with fewer numbers than was observed for non-wrinkled bilayers.

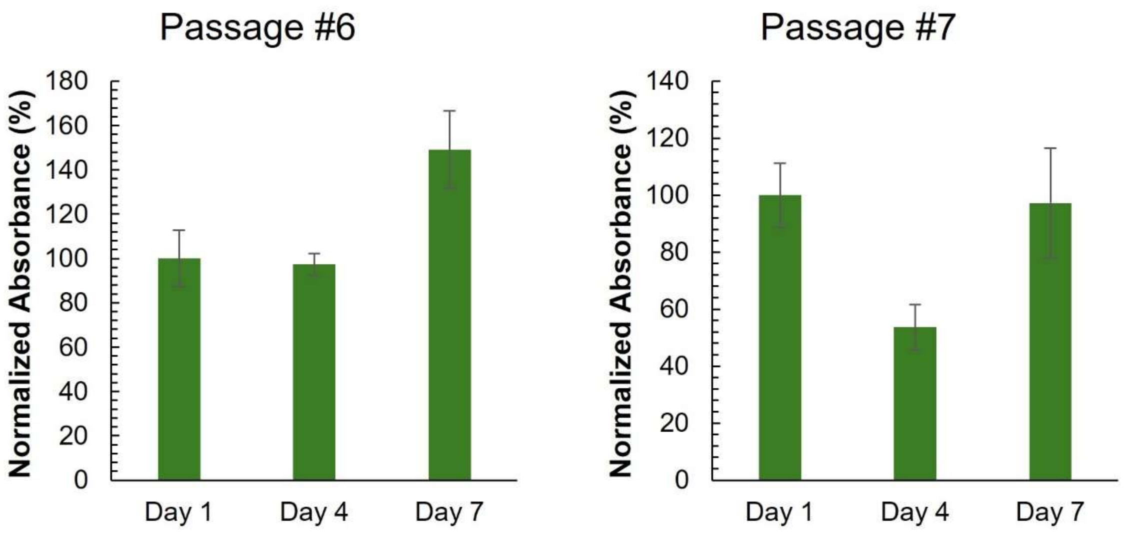


**Figure S5. MTT assay of HEKs cultured on nanofibrous membranes.** Although membranes appear to support the growth of cells, as indicated by an increase in metabolic activity over time, HEK passage number significantly affects proliferative capacity. For passage #7, a notable decrease in metabolic activity was observed on day 4 compared to day 1. *n = 3*.

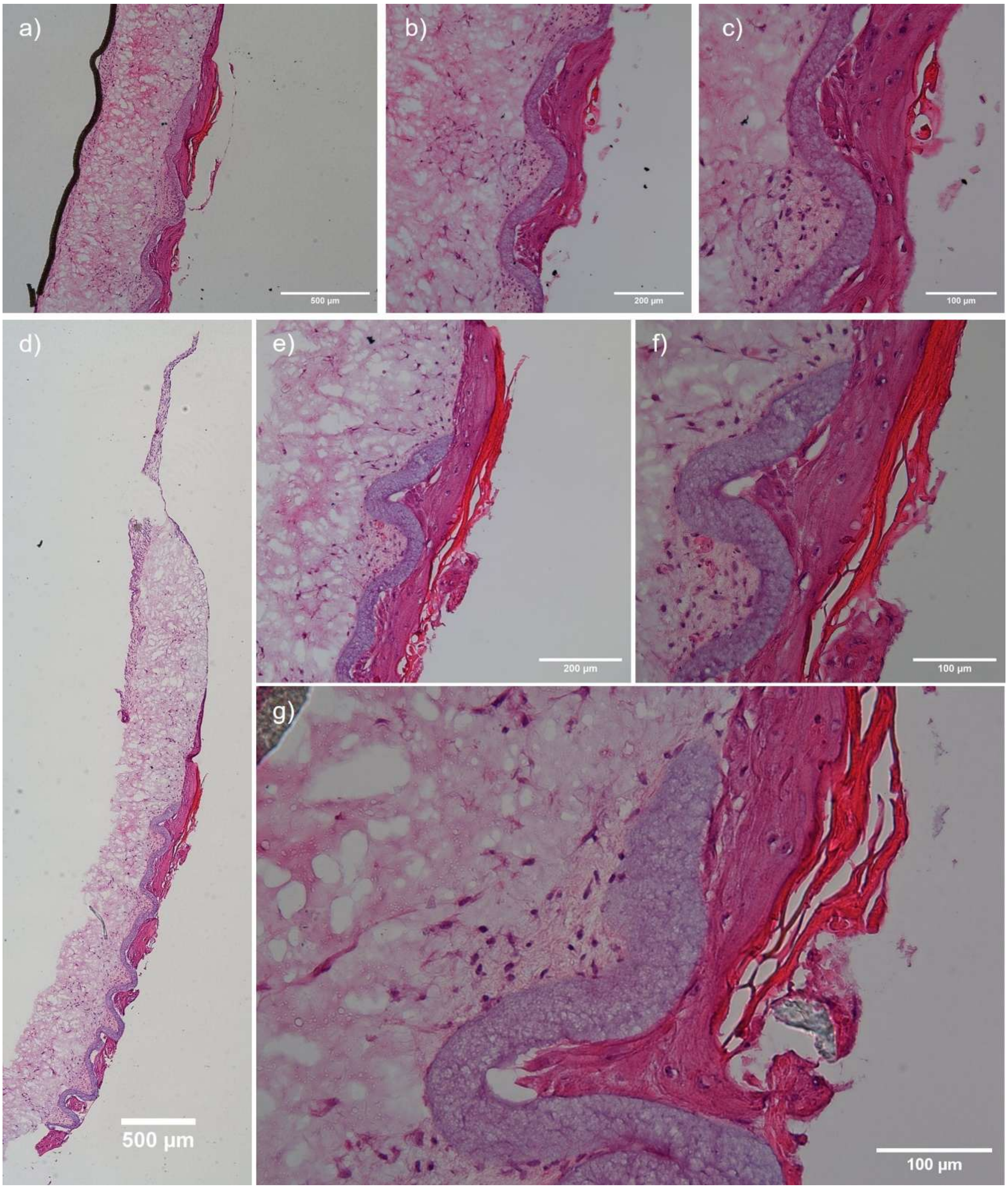


**Figure S6. Histological sections of bilayer skin models using an improved experimental setup.** H&E staining reveals considerably thicker epithelial layers upon allowing confluent keratinocyte monolayer formation prior to initiating wrinkling at the beginning of the co-culture period. Cornified epithelial layers are stained bright red.
